# DNA barcoding–based paired daughter cell analysis reveals division preferences of hematopoietic stem cells

**DOI:** 10.64898/2026.07.24.739718

**Authors:** Tsuyoshi Fukushima, Akira Nishiyama, Shuhei Koide, Tomoya Isobe, Tomohiro Yabushita, Shuhei Asada, Susumu Goyama, Atsushi Iwama, Satoshi Yamazaki, Tomohiko Tamura, Toshio Kitamura, Toshio Suda, Yosuke Tanaka

## Abstract

Hematopoietic stem cells(HSCs) maintain their pools by stem-stem division and produce mature blood cells through stem-progenitor or progenitor-progenitor division. A paired daughter cell(PDC) assay combined with single cell transplantation is a powerful method to compare the lineage outputs of two HSC daughter cells. However, single-cell transplantation precludes large-scale analysis of daughter-cell pairs, as only one cell can be transplanted per recipient. Here, we developed a DNA barcoding–based PDC assay to overcome this limitation, enabling simultaneous analysis of 476 daughter pairs from individual HSC divisions and revealing that daughter-cell fates are coordinated and that HSC division patterns are biased toward stem–stem and progenitor–progenitor divisions rather than stem–progenitor divisions. These findings indicate that HSC fate outcomes are directed toward symmetric division outcomes. Integration of single-cell RNA sequencing with DNA barcoding revealed a continuum of HSC states—from balanced HSCs to myeloid-biased HSCs and ultimately to a low-output HSC subset—in which progressively reduced production of mature blood cells relative to stem cell expansion, a proxy for stem–stem division bias, exhibits distinct activities of transcription factors and signaling pathways. Overall, our analysis uncovers characteristic patterns of HSC division and links stem maintenance with distinct molecular features.

## Introduction

Hematopoietic stem cells (HSCs) sustain hematopoiesis throughout life by both maintaining the stem cell pool and producing mature blood cells(Ito et al., 2016; Seita and Weissman, 2010; Yamamoto et al., 2013). To support these two functions, HSCs are thought to undergo three distinct types of cell division: stem–stem, stem–progenitor, and progenitor–progenitor divisions. Stem–stem divisions expand the stem cell pool. Stem–progenitor divisions maintain stem cells while generating differentiated progeny. Progenitor–progenitor divisions are thought to sacrifice stem cell properties to produce differentiated cells(Ito *et al*., 2016; Seita and Weissman, 2010; Yamamoto *et al*., 2013). Paired daughter cell (PDC) assays enable the analysis of two daughter cells derived from a single mother HSC(Suda et al., 1984), allowing us to investigate HSC division patterns. Previous studies using paired daughter cell combined with single cell transplantations have analyzed serval dozen pairs of HSC division(Ito *et al*., 2016; Yamamoto *et al*., 2013). However, it remains difficult to obtain a comprehensive picture of HSC division dynamics, as analyzing a large number of daughter cell pairs poses enormous challenge. This is because the conventional methods require each daughter cell derived from a single mother HSC to be transplanted into a separate recipient mouse—a technically demanding procedure that requires numerous recipient animals and has limited efficiency.

DNA barcoding assays allow tracing the lineage output of single HSCs in large numbers at once(Naik et al., 2013; Rodriguez-Fraticelli et al., 2020; Rodriguez-Fraticelli et al., 2018; Sun et al., 2014). However, daughter cells derived from the same single cell share the identical barcode, making it impossible to distinguish the tracing outcomes of individual daughter cells. In this study, we develop a method for lineage tracing of a large number of paired daughter cells from HSCs using DNA barcoding, enabling the simultaneous analysis of 476 HSC divisions. In addition, the combination of single cell RNA sequence (scRNA-seq) and DNA barcoding has revealed the molecular features of HSCs that produce distinct outputs.

## Results

### DNA barcodes allow for a large number of paired daughter cell assays

We have developed a system that uses DNA barcodes to assess the lineage output of paired daughter cells of HSCs. The method is as follows: (1) HSCs were isolated from mouse bone marrow (Figure. 1A and . S1A). (2) The isolated HSCs were transduced with DNA barcodes using lentiviral vectors carrying a DNA barcode (Figures 1A, S1B, and S1C). (3) The transduced HSCs were cultured until the first cell division (Figure 1A). (4) The cultured cells were then split into two halves (Figure 1A). (5) Each of the two halves was transplanted into a separate recipient mouse (Figure 1A). (6) The DNA barcodes were recovered from differentiated progeny of each lineage and analyzed by next-generation sequencing (Figures 1B, and S1D-S1F). This approach enables the identification of division patterns of individual HSCs during in vitro culture by functionally evaluating the long-term repopulating and lineage output potential of each daughter cell following transplantation. After undergoing a single division, barcoded HSCs are split into two halves, and the probability that a given daughter pair is separated into different halves is theoretically 50%. It is expected that the barcodes of the separated daughter pairs will be detectable in both recipient mice after transplantation. Such barcodes are considered to originate from the same daughter pair (Figure 1C). To determine the optimal culture duration in which HSCs undergo a single division but rarely divide twice before transplantation, live-cell imaging was performed. At 60hrs culture, 92.5% of cells had undergone at least one division, whereas only 17.2% had undergone two divisions (Figure 1D), indicating that 60 hours is the optimal culture time (Figure 1D). Furthermore, analysis of DNA barcodes from HSCs that were cultured for different periods of time and then divided in half revealed that the same barcode appeared in both wells when cells were divided after 60 hours of culture, but not after 36 hours of culture (Figure 1E). Based on these results, the culture duration prior to transplantation was set to 60 hours.

**Figure 1.**
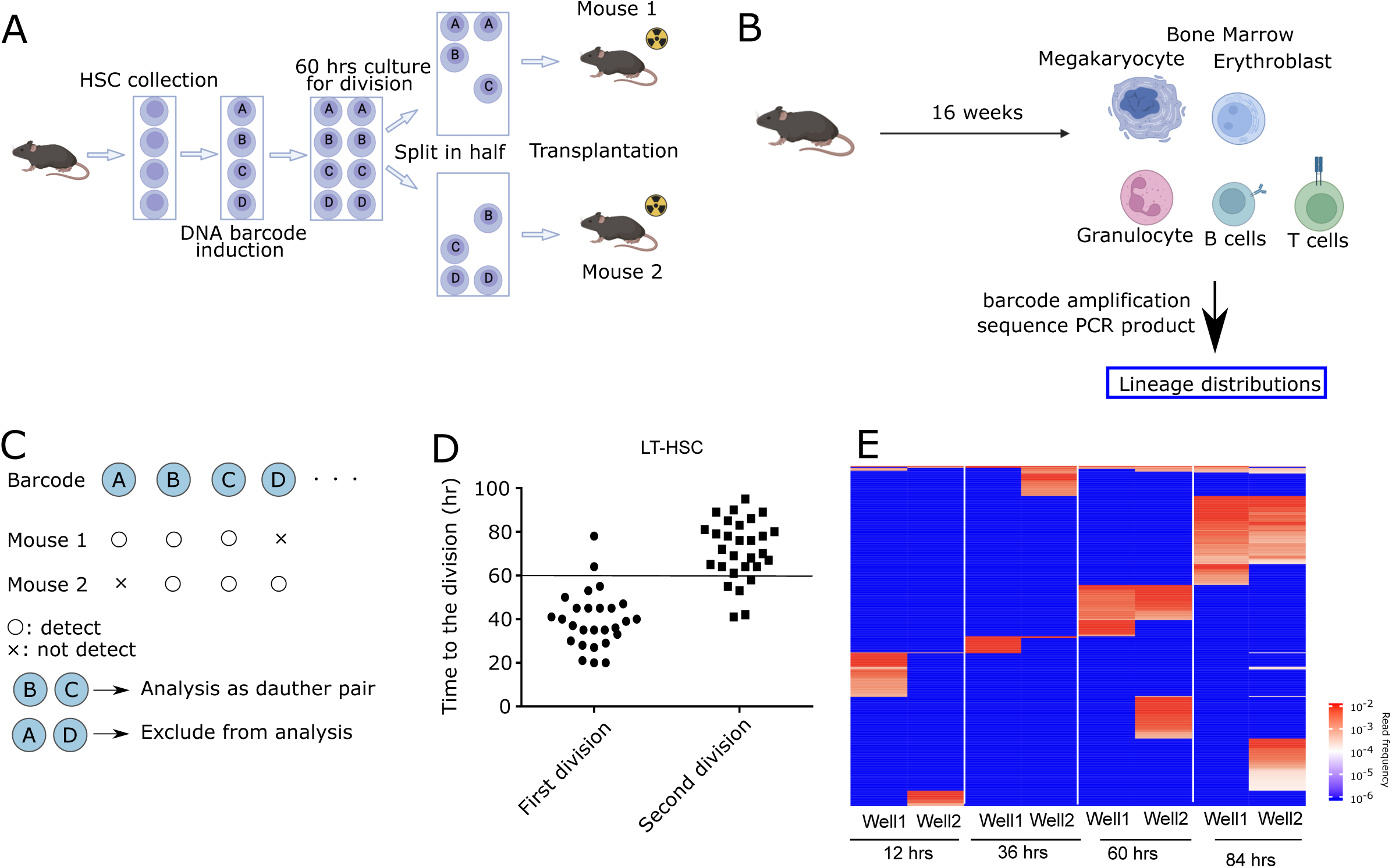
DNA barcoding–based paired daughter cell (PDC) assay enables large-scale analysis of HSC division patterns. **A**, Schematic overview of the DNA barcoding–based paired daughter cell assay. Hematopoietic stem cells (HSCs) were isolated from mouse bone marrow and transduced with lentiviral vectors carrying unique DNA barcodes. After culture for 60 hours to allow the first cell division, cells were randomly split into two equal fractions, and each fraction was transplanted into a separate recipient mouse. **B**, Experimental design for lineage output analysis. Sixteen weeks after transplantation, mature hematopoietic populations—including megakaryocytes (Mk), erythroblasts (Er), granulocytes/Monocyte (GM), B cells, and T cells—were isolated from bone marrow. DNA barcodes were amplified by PCR and analyzed by next-generation sequencing to determine lineage distributions. **C**, Strategy for identifying paired daughter clones. Barcodes detected in both recipient mice were considered to originate from a separated daughter pair derived from a single HSC division. Barcodes detected in only one recipient were excluded from paired analysis. **D**, Live-cell imaging analysis of long-term HSCs (LT-HSCs) showing time to first and second divisions. Most cells underwent the first division within approximately 60 hours, whereas second divisions occurred later. **E**, Heatmap showing barcode read frequency in split wells after different culture durations (12, 36, 60, and 84 hours). Identical barcodes were detected in both wells after 60 hours, indicating successful separation of daughter pairs. Color scale represents read frequency.

### DNA barcoding–based paired daughter cell analysis reveals division preferences of HSCs

To analyze HSC division patterns, barcodes were retrieved from megakaryocyte progenitors (Mk), erythroid progenitors (Er), granulocytes/monocytes (GM), B cells, and T cells in the bone marrow of mice 16 weeks after transplantation using the aforementioned (Figures 1A-1B and S1A-S1G). Because barcodes repeatedly detected across independent experiments are more likely to be independently assigned to multiple cells, they pose a particular risk in paired daughter assays by potentially causing unrelated cells to be misidentified as daughter-cell pairs. Therefore, barcodes observed in at least two of three independent experiments were excluded from further analysis (Figure S2A). A total of 3,542 daughter clones were detected, comprising the 952 daughter cells (26.8% of all daughter clones) derived from 476 daughter pairs (Figures S2A-C). Unsupervised k-means clustering classified the daughter cells into nine distinct clusters (Figure 2A). It has been reported that long-term engrafting HSCs produce either all lineages after 16 weeks in a primary transplantation, or that they lack T cells, or both B and T cells (Carrelha et al., 2018; Yamamoto et al., 2018). In other words, HSCs are defined as cells that produce all megakaryocyte, erythroid, and granulocyte/monocyte lineages, while progenitors defined as the cells whose produced cells do not contain one of these lineages. Therefore, in this study, the daughter cells that generate all lineages 16 weeks after transplantation are referred to as balanced HSCs (Bal-HSCs), whereas the daughter cells that produce megakaryocyte, erythroid, and granulocyte/monocyte lineages (Mk-Er-GM) or Mk-Er-GM-B lineages are referred to as myeloid-HSCs (My-HSCs) (Figures 2A and 2B). The daughter cells in other clusters are referred to as GBT, MyB, Mk, Er, GM, B, and T progenitors (progs) according to the lineage of the cells produced (Figures 2A, 2B and S2D). The relationships between 476 daughter pairs were analyzed. Of the 476 daughter pairs, 50 underwent stem–stem division, 34 underwent stem–progenitor division, and 392 underwent progenitor–progenitor division (Figure 2C). Pairs between stem cells (Bal-HSCs and My-HSCs) were observed more frequently than expected, whereas pairs between stem cells and progenitors (GBT, MyB, Mk, Er, GM, B, and T progs) occurred significantly less frequent than would be expected from random pairing, and pairs between progenitors occurred at frequencies consistent with random expectations (Figures 2C and 2D). Although the absolute frequencies of stem–stem and stem–progenitor divisions may depend on the biological context, our results indicate that HSC fate outcomes are not randomly assigned. Instead, stem cell divisions show a tendency toward symmetric outcomes as stem–stem and progenitor–progenitor divisions occurring rather than asymmetric outcomes as stem–progenitor divisions. Because our primary analysis focused on clones that engrafted in both recipient mice, paired clones in which one daughter failed to achieve long-term engraftment or remained below the detection limit may have been underestimated. To address this potential bias, we performed an additional analysis assuming that the counterpart of each clone detected in only one recipient represented a progenitor clone that either lacked long-term repopulating capacity or remained below the detection limit. Under this assumption, the same overall statistical trend was observed (Figure S2E), indicating that our conclusions are robust to this potential source of bias. Analysis by cell type revealed that divisions between Bal-HSCs, as well as between Bal-HSCs and My-HSCs, were significantly enriched (Figure 2E). In contrast, no divisions between My-HSCs were observed (Figure 2E). However, since only a small number of My-HSCs were detected, it could not be determined whether divisions between My-HSCs are genuinely rare. Furthermore, pairs between Bal-HSCs and progenitors—except for GBT progenitors—were significantly underrepresented (Figure 2E), indicating that Bal-HSCs are less likely to undergo stem–progenitor divisions. In progenitor–progenitor divisions, enrichment was observed for certain pairs of the same cell type, such as MyB–MyB, B–B, and GM–GM (Figure 2E). Together, DNA barcoding–based paired daughter cell analysis revealed that HSCs preferentially undergo stem–stem or progenitor–progenitor divisions rather than stem–progenitor divisions, highlighting the division preferences of HSCs.

**Figure 2.**
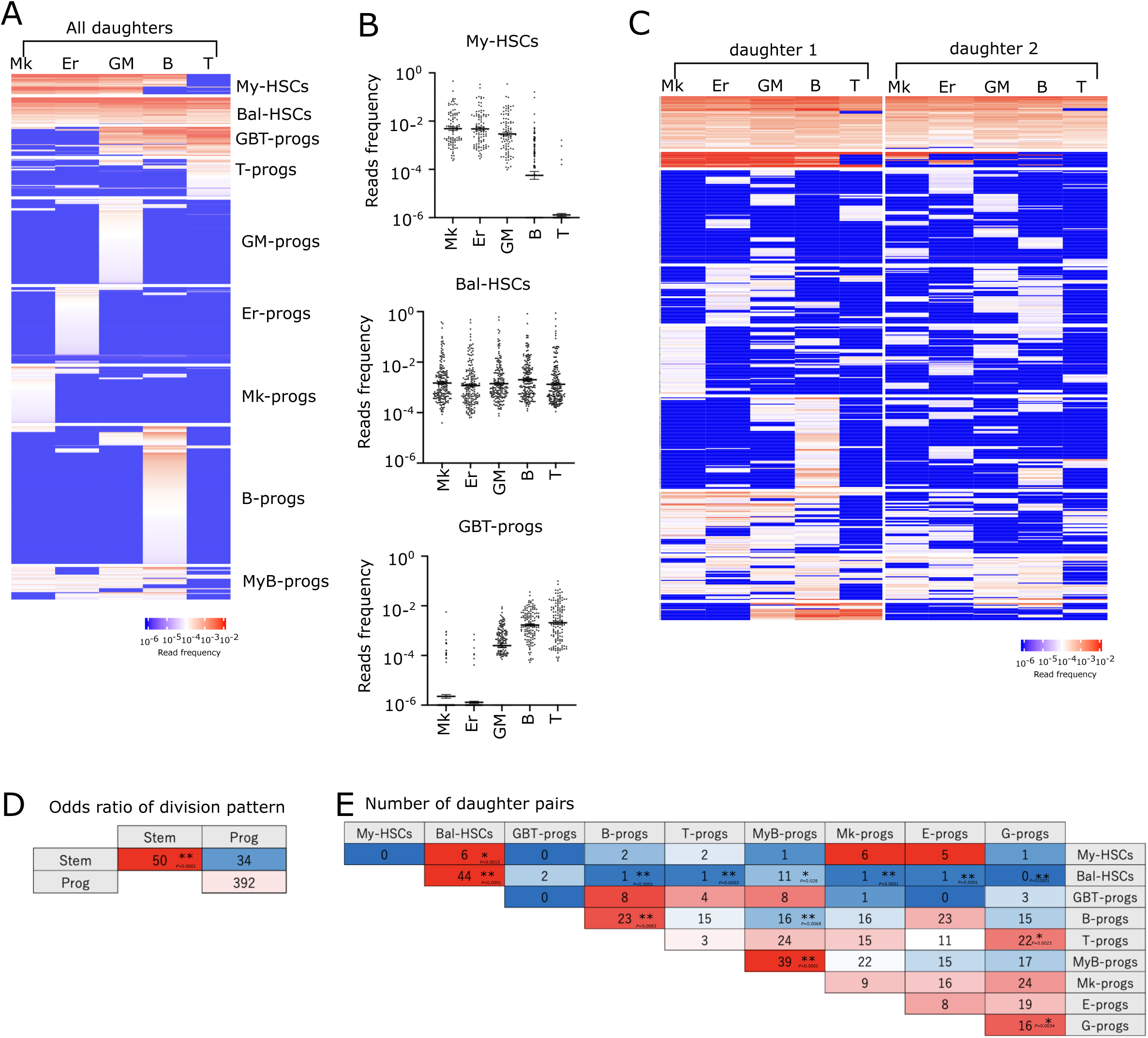
Large-scale paired daughter cell analysis reveals preferential stem–stem and progenitor–progenitor divisions of HSCs. **A**, Heatmap showing lineage output profiles of all detected daughter clones (n = 3,477). Each row represents a daughter clone, and columns indicate normalized barcode read frequencies in megakaryocyte (Mk), erythroid (Er), granulocyte/monocyte (GM), B cell (B), and T cell (T) compartments 16 weeks after transplantation. Unsupervised k-means clustering classified daughter clones into nine distinct clusters: balanced HSCs (Bal-HSCs), myeloid-biased HSCs (My-HSCs), GBT progenitors (GBT-progs), T progenitors (T-progs), GM progenitors (GM-progs), Er progenitors (Er-progs), Mk progenitors (Mk-progs), B progenitors (B-progs), and MyB progenitors (MyB-progs). **B**, Dot plot showing the lineage output of My-HSCs, Bal-HSCs, and GBT-progs. Each dot represents a daughter clone; y-axis indicates barcode read frequency (log scale). **C**, Heatmaps of lineage outputs for paired daughter cells (n = 476 daughter pairs), shown separately for daughter 1 and daughter 2. Rows correspond to paired clones derived from a single HSC division. **D**, Table showing the odds ratios comparing observed versus expected frequencies of stem–stem, stem–progenitor, and progenitor–progenitor pairings under random pairing assumptions. Statistical significance was assessed using bimodal test (p values indicated). **E**, Table showing the number of daughter pairs for each combination of cell types. Color intensity reflects odds ratio relative to random expectation. Statistical significance was assessed using bimodal test (p values indicated). *p < 0.05 and **p < 0.01. n.s., not significant.

### Low-output HSCs preferentially undergo stem–stem divisions with high stemness programs

To identify the molecular signature of each cell types, we combined random DNA barcoding and scRNA-seq analyses. The barcode is transcribed with BFP and can be detected by scRNA-seq (Figure S1B)(Adamson et al., 2016), allowing the clonal identity of each cell profiled by scRNA-seq to be determined. This information can be integrated with lineage output data of the corresponding clones obtained from mature blood cells. To link lineage output characteristics with the molecular features of HSCs, immunophenotypic HSCs were isolated from the bone marrow of mice 16 weeks after transplantation performed using the strategy shown in Figures 1A and 1B, and subjected to scRNA-seq. After filtering, barcodes were detected in 14,431 of 15,642 cells (92.2%). A total of 368 daughter clones were detected in scRNA-seq dataset. Bal-HSCs and My-HSCs are capable of long-term HSC maintenance; both Bal-HSCs and My-HSCs contributed to the pool of immunophenotypic HSCs in the bone marrow 16 weeks after transplantation (Figure. 3A). This supports the classification of these fractions as HSCs, as shown in Figure. 2. Furthermore, the lineage-biased HSC fractions identified in our study are highly consistent with those reported previously using single-cell transplantation (Carrelha *et al*., 2018; Yamamoto *et al*., 2018) without in vitro culture or lentiviral barcoding. This similarity indicates that these HSC populations can be reliably observed even after in vitro culture and lentiviral transduction, suggesting that the culture and barcoding procedures did not substantially alter their functional properties. In addition, we identified a subset of clones that contributed little to differentiated cells during the primary transplantation and defined them as low-output HSCs (Low-HSCs) (Figure 3A). The Low-HSC population classified in this study corresponded to previously described subset (Rodriguez-Fraticelli *et al*., 2020). Low-HSCs exhibited a lower mature blood cell output–to–stem expansion ratio compared with other HSC subsets (Figure 3B), suggesting that this subset preferentially undergoes stem–stem divisions. Consistently, analysis of paired daughter cell data showed that 4 out of 6 divisions occurred between two Low-HSCs, and 1 out of 6 involved Low-HSCs and another stem cell, further supporting the predominance of stem–stem divisions in this subset (Figure 3C). Next, to molecularly classify single cells, dimensionality reduction was performed using Uniform Manifold Approximation and Projection (UMAP)(Becht et al., 2018), followed by clustering (Figure 3D). Cell type annotation based on the ImmGen(Heng and Painter, 2008) reference dataset classified Clusters 1–3 as HSCs (HSC1-3), whereas Clusters 4 and 5 were annotated as Megakaryocyte–Erythroid Progenitors (MEP) and Early T cell Progenitors (ETP), respectively. (Figures 3E and S3A-E). Among HSC1–3, HSC1 showed higher expression of stem cell–associated genes, including Mllt3, Hlf, and Mecom, and exhibited the highest stem gene scores(Hamey and Göttgens, 2019; Wilson et al., 2015) (Figures 3E and 3F). In contrast, HSC2 and HSC3 expressed higher levels of genes associated with more differentiated HSC states, such as Cd34 and Gata1, and displayed lower stem gene scores (Figures 3E and 3F). HSC3 also exhibited elevated cell cycle scores, with the majority of cells being classified in the G2/M phase (Figure S3F). Furthermore, single cells classified as Low-HSCs were preferentially enriched in HSC1 (Figure. 3G and . S3G), suggesting that Low-HSCs tend to maintain a state of high stem gene expression. These findings further support the notion that Low-HSCs predominantly undergo stem–stem divisions. Collectively, these results establish Low-HSCs as a subset that preferentially undergoes stem–stem divisions.

**Figure 3.**
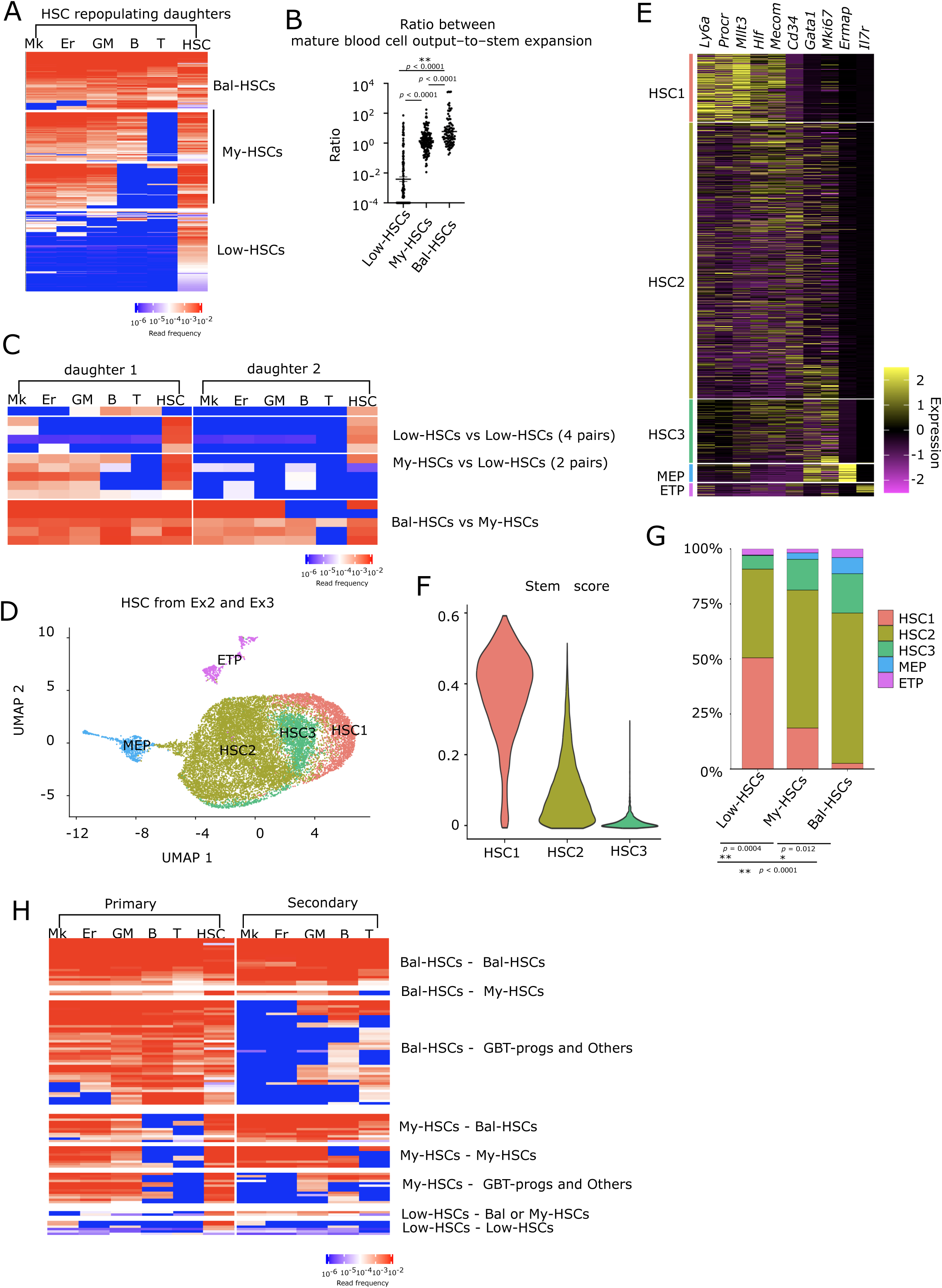
scRNA-seq identifies a Low-HSCsubset preferentially undergoing stem–stem divisions and characterized by high stemness programs. **A**, Heatmap showing lineage outputs and stem cell contribution of HSC-repopulating daughter clones 16 weeks after transplantation. Each row represents a clone classified as Bal-HSCs, My-HSCs, or Low-HSCs. Columns indicate normalized barcode read frequencies in mature lineages (Mk, Er, GM, B, T) and the HSC compartment. **B**, Ratio of mature blood cell output to stem expansion for each HSC subset. The y-axis represents the ratio between total mature lineage output and HSC contribution per clone. Statistical significance was assessed by one-way ANOVA followed by Tukey’s multiple comparisons test (p values indicated). **C**, Lineage output heatmaps of representative paired daughter clones, grouped by combination type (Low-HSCs vs Low-HSCs, My-HSCs vs Low-HSCs, Bal-HSCs vs My-HSCs). Rows correspond to paired daughters (daughter 1 and daughter 2). Low-HSC pairs show low mature lineage contribution in both daughters while retaining HSC output. **D**, UMAP visualization of single-cell transcriptomes derived from immunophenotypic HSCs (CD150⁺CD48^-^c-Kit⁺ Sca-1⁺ Lin⁻ from Ex2 and CD150⁺ c-Kit⁺ Sca-1⁺ Lin⁻ from Ex3) isolated 16 weeks after transplantation. Unsupervised graph-based clustering using the Seurat pipeline identified 10 clusters (c1–c10). **E**, Heatmap of representative marker genes across clusters. Rows represent individual cells and columns represent genes. **F**, Stemness score distribution across clusters 1–3. **G**, Proportion of transcriptional clusters across Bal-HSCs, My-HSCs, and Low-HSCs. Statistical significance was assessed using pairwise Wilcoxon rank-sum tests with Benjamini–Hochberg correction for multiple comparisons (p values indicated). **H**, Lineage output heatmaps of paired daughter clones following primary and secondary transplantation (n = 281 clones). Columns represent normalized barcode read frequencies in Mk, Er, GM, B, T, and HSC compartments. Each raw represent daughter clone. *p < 0.05 and **p < 0.01. n.s., not significant.

Secondary transplantation revealed transitions from My-HSCs to Bal-HSCs (Figure 3H), consistent with a previous report (Yamamoto *et al*., 2018). In addition, Bal-HSCs tended to lose the ability to produce myeloid cells such as MK, RBC and GM lineages during the secondary transplantation (Figure 3H), consistent with prior studies showing that myeloid chimerism declines earlier than lymphoid chimerism (Yamamoto *et al*., 2013; Yamamoto *et al*., 2018). Interestingly, 2 of 8 generated My-HSC and Balanced-HSC-like outputs (Figure 3H). These subsets resemble previously described latent-HSCs(Morita et al., 2010). In contrast, 6 of 8 Low-HSCs contribute minimally to hematopoiesis even after secondary transplantation. Thus, at least under the regenerative stress associated with secondary transplantation, these cells did not appear to play a significant role in mature blood cell production.

### Transcriptional programs of Low-HSCs associated with preferential stem–stem divisions

Finally, we sought to identify candidate genes regulating stem–stem divisions by analyzing the molecular signature of Low-HSCs. Low-HSCs showed higher expression of Mllt3 which is essential for long-term bone marrow reconstitution (Calvanese et al., 2019). Additionally, Vwf, a marker gene of My-HSCs (Sanjuan-Pla et al., 2013) were expressed at higher levels in My-HSCs than in Bal-HSCs and at even higher level in Low-HSCs (Figures 4A and S4A). In contrast, Runx3(Meng et al., 2023), which has been reported to be more active in Bal-HSCs than in My-HSCs, showed lower expression in Low-HSCs than in My-HSCs (Figures 4A and S4A). These enriched or depleted genes in Low-HSCs also include surface markers such as Neo1(Gulati et al., 2019), Cd74, and Cd52 (Figures 4A and S4A). Although the expression of these markers changes continuously across balanced-HSCs, My-HSCs, and Low-HSCs, their individual expression levels or combinations may contribute to the development of future FACS-based strategies for the prospective enrichment and isolation of functionally distinct HSC states. Furthermore, pathway analysis–based inference of transcription factor activity identified Pparg and Mef2c as potentially active transcription factors in Low-HSCs (Figure 4B). Pparg has been previously reported to promote stem–stem divisions upon activation (Ito *et al*., 2016). We have also shown that HSCs with high cytosolic calcium exhibit enhanced engraftment potential and dormancy (Fukushima et al., 2019), making it particularly interesting that Mef2c, calcium-dependent signaling (Canté-Barrett et al., 2014), was predicted as one of the transcription factors highly active in Low-HSCs in our analysis. Analysis of signaling pathway activity further showed that pathways known to regulate HSC maintenance, including WNT (Sugimura et al., 2012) and hypoxia (Takubo et al., 2010) signaling, were elevated in Low-HSCs (Figure 4C). In contrast, candidate transcription factors with low activity in Low-HSCs included myeloid-related factors such as Spi1, Cebpa, and Irf4, as well as inflammation-associated factors such as Nfkb2 and Nr4a1 (Figure 4B). Signaling pathway activity inference indicated reduced TNFα and MAPK signaling in Low-HSCs (Figure 4C). Inflammation is known to impair HSC maintenance and promote differentiation (Takizawa et al., 2017), suggesting that inflammatory responses may also hinder stem–stem divisions. Collectively, these results define Low-HSCs as a subset that preferentially undergoes stem–stem divisions and their transcriptional signatures, providing insight into the molecular mechanisms underlying HSC self-renewal.

**Figure 4.**
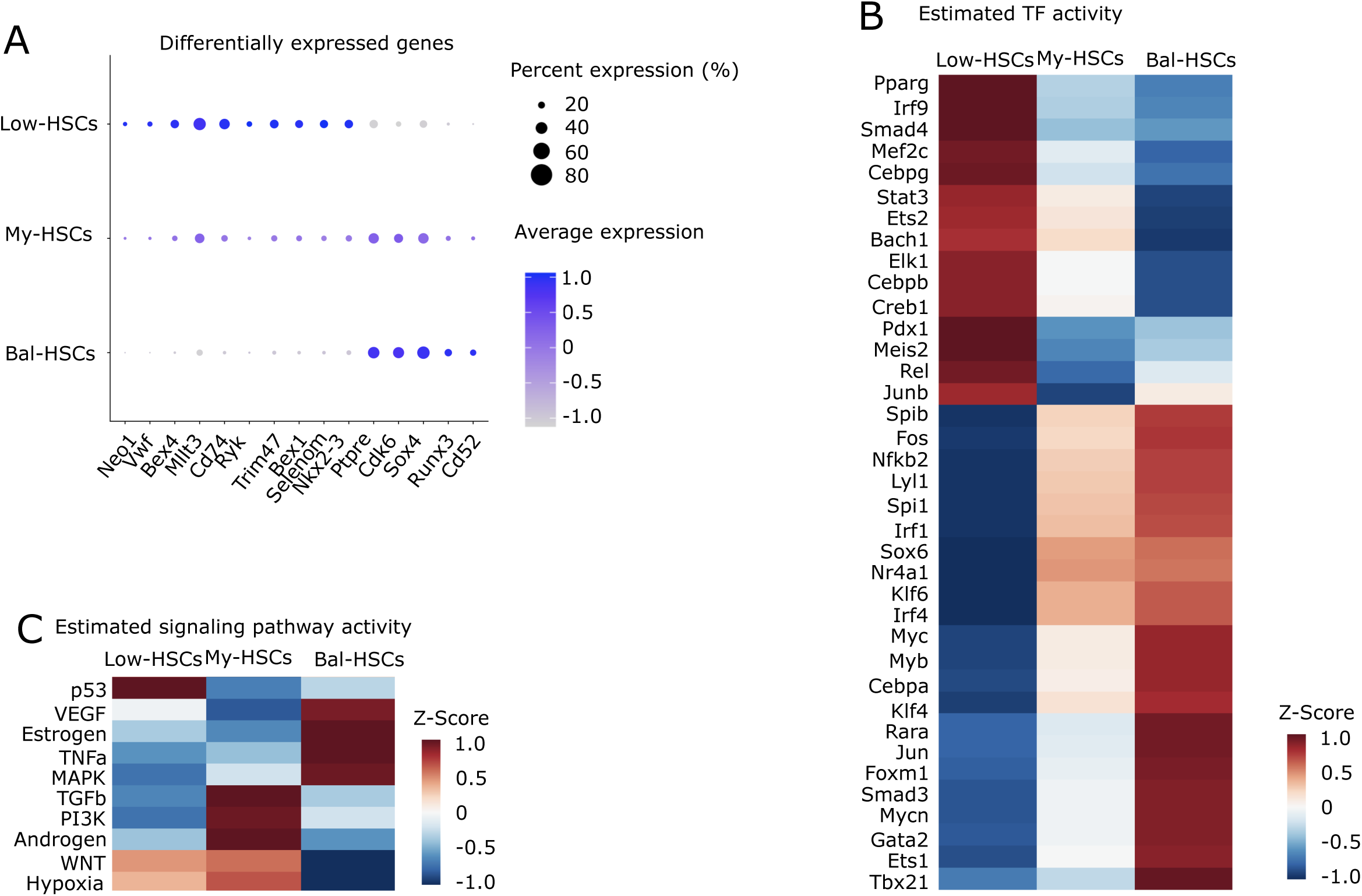
Transcriptional programs and candidate regulators associated with stem–stem division–biased Low-HSCs. **A**, Dot plot showing differentially expressed genes across Low-HSCs, My-HSCs, and Bal-HSCs. Dot size represents the percentage of cells expressing each gene, and color intensity indicates average expression level. **B**, Heatmap of inferred transcription factor (TF) activity estimated using decoupleR evaluated using cells in HSC1-3. Rows represent individual TFs and columns represent HSC subsets (Low-HSCs, My-HSCs, and Bal-HSCs). Color intensity indicates Z-scored TF activity across subsets. **C**, Heatmap of inferred signaling pathway activity estimated using decoupleR evaluated using cells in clusters 1. Rows represent individual signaling pathway and columns represent HSC subsets (Low-HSCs, My-HSCs, and Bal-HSCs). Color intensity indicates Z-scored TF activity across subsets.

## Discussion

In this study, we enabled large-scale PDC analysis of 476 daughter pairs and showed that HSCs rarely undergo stem–progenitor divisions, instead preferentially executing stem–stem divisions. In conventional paired daughter cell assays combined with single-cell transplantation, the number of analyzable pairs has been limited to several dozen, thereby precluding statistically robust analyses of HSC division patterns (Ito *et al*., 2016; Yamamoto *et al*., 2013).

Compared with conventional paired daughter cell assays based on single-cell transplantation(Ito *et al*., 2016; Yamamoto *et al*., 2013), the barcode-based method used in this study likely provides higher sensitivity for detecting progenitors which fail to sustain long-term multi lineage contribution after transplantation (Figure 2A). Despite this increased sensitivity for progenitor detection, stem–progenitor divisions remained limited, whereas stem–stem and progenitor–progenitor divisions were preferentially enriched. These findings indicate that our approach captures characteristic features of HSC division patterns and provides a useful framework for comprehensively evaluating HSC division dynamics.

CP-tracer, which relies on the accumulation of randomly acquired mutations to trace the progeny of individual HSCs, has revealed the long-term plasticity of HSC fate(Fukushima et al., 2025). However, because the timing of mutation acquisition is stochastic, the initial cell division and its immediate progeny cannot be directly and precisely defined. In contrast, by experimentally separating daughter pairs generated from the first division of an HSC and subsequently tracking their functional fates, the present study enables a more direct assessment of the fate relationship between the first-generation daughter cells arising from the same parental HSC.

Interestingly, whereas a minority of Low-HSC-derived clones generated My-HSC- or Balanced-HSC-like outputs, the majority maintained a Low-HSC pattern even after secondary transplantation. An important unresolved question is whether Low-HSCs represent a functionally competent reserve stem-cell pool that can contribute to hematopoiesis under specific conditions, or alternatively a low-output HSC state in which a strong bias toward stem–stem division results in persistently limited mature blood-cell production. Aging and clonal hematopoiesis are associated with the myeloid bias and knock-out of Dnmt3a which is known to driver of clonal hematopoiesis enhance stem-maintenance-bias. These observations raise the possibility that an extreme stem-maintenance-biased state could contribute to pathological hematopoiesis by favoring long-term stem-cell persistence while limiting productive blood-cell output.

Stem cell factor (SCF) and thrombopoietin (TPO) are well-established regulators of HSC maintenance and self-renewal both in vivo and ex vivo.(Ding et al., 2012; Nakauchi et al., 2001; Qian et al., 2007; Wilkinson et al., 2019). Even within SCF/TPO-based culture systems, alterations in cytokine concentrations can substantially affect HSC expansion, maintenance of long-term repopulating activity, and lineage commitment(Wilkinson *et al*., 2019). In addition, alternative cytokine combinations or the use of different culture system may influence the divison pattern. The in vitro culture environment does not fully recapitulate the native bone marrow niche. A definitive understanding of HSC fate decisions would ideally require direct identification and functional assessment of the two daughter cells generated from a single HSC division in vivo. However, tracking and distinguishing these daughter cells within the native bone marrow environment remains technically extremely challenging. Future methodological advances enabling in vivo discrimination of daughter HSC pairs will therefore be important for further elucidating the mechanisms governing HSC fate decisions.

In summary, this work establishes a scalable and integrative approach that connects stem cell division patterns with underlying molecular programs, thereby enabling a more systematic dissection of the principles governing stem cell fate decisions.

## Acknowledgments

We thank the Flow Cytometry Core at The Institute of Medical Science, The University of Tokyo for their help. This project was supported by JSPS KAKENHI Grant Number JP24K11520 and a grant from The Uehara Memorial Foundation, SENSHIN Medical Research Foundation to Y.T., ‘‘Grant-in-Aid for Challenging Exploratory Research’’ from JSPS Grant Number 20K21613 and the Japan Science and Technology Agency ACT-X program Grant Number JPMJAX25L9 to Tsu.F. and MEXT Promotion of Distinctive Joint Usage/Research Center Support Program Grant Number JPMXP0618217493 at the Advanced Medical Research Center, Yokohama City University.

## Author contributions

Tsu.F. and Y.T. designed most of the experiments. Tsu.F. and Y.T. performed most of the experiments. A.N and S.K. performed single cell RNA-seq. T.Y. supported to DNA-seq of barcode. A.N. and T.I. supported single cell RNA-seq data analysis. Tsu.F., Y.T., S.Y., S.G., S.A. and T.K. wrote the manuscript. Y.T, T.K., T.T., S.T., S.Y., S.G. and A.I. supervised and coordinated the project.

## Data and Material Accessibility

Materials are available, subject to material transfer agreement requests submitted to Y.T.. scRNA-sequencing data have been deposited at GEO “GSE321678” and are publicly available as of the date of publication. All other data available in the manuscript or supplementary materials are available from corresponding author upon reasonable request.

## Method

### Mice

Female C57BL/6-Ly5.2 (Ly5.2) were purchased from Japan SLC (Shizuoka, Japan). Eight- to twelve-week-old male and female mice served for experiments unless otherwise noted. All mice were housed at 22 ±2 ℃ and 12:12 hrs light: dark cycle. The experiments were approved by the Committee on the Ethics of Animal Experiments, University of Tokyo.

### Isolation of Hematopoietic Stem Cells

BM hematopoietic cells were isolated from femurs and tibias, humerus, sternum, pelvic and vertebra by clashing and depleted of red blood cells by ACK Lysing Buffer (Thermo Fisher Scientific). Bone marrow cells were incubated with Lineage antibodies (1:1000) (Anti-mouse CD5-biotin Antibody (clone 53-7.3) (BioLegend, 100604), Anti-mouse/human CD45R/B220-biotin Antibody (clone RA3-6B2) (BioLegend, 103204), Anti-mouse TER-119-biotin Antibody (clone TER-119) (BioLegend, 116204), Anti-mouse Gr-1-biotin Antibody (clone RB6-8C5) (BioLegend, 108404), Anti-mouse/human CD11b-biotin Antibody (clone M1/70) (BioLegend, 101204)) in 1 mL for 10 minutes at 4 ℃. After adding 45 mL of PBS and centrifuging for 5 minutes, the supernatant was discarded. This process was repeated twice. The cells were then incubated with 10 mL of PBS and 90 μL of SA microbeads (Miltenyi Biotec, 130-090-858) for 15 minutes at 4 ℃. After adding 45 mL of PBS and centrifuging for 5 minutes, the supernatant was discarded. This process was repeated twice. Negative selection was then performed using LS columns (Miltenyi Biotec, 130-042-401). The cells from the negative fraction were centrifuged for 5 minutes, and the supernatant was removed. The cells were incubated with a cocktail of antibodies, including CD150-PE (clone TC15-12F12.2) (1:200), CD48-APC-cy7 (clone HAM48-1) (1:200), c-Kit-PEcy7 (clone 2B8) (1:200), Sca-1-APC (clone D7) (1:400) and Streptavidin-BV605(1:400) in 400µl for 15 minutes at 4 ℃. Following the addition of 1 mL of PBS and centrifugation for 5 minutes, the supernatant was discarded. The cells were resuspended in 1 mL of PBS and sorted for the CD150^+^CD48^-^Lin^-^cKit^+^Sca1^+^ fraction using FACS Aria with single stain control.

### Plasmid Preparation

Electroporation was performed by adding 1µl of the plasmid Perturb-Seq Guide Barcodes (GBC) Library (Addgene Plasmid # #85968) (diversity > 100,000) (Adamson *et al*., 2016) to 100 µl of MegaX DH10B competent cells (Invitrogen, C640003). The mixture was subjected to electroporation at 2.0 kV, 200 Ω, and 25 µF. Immediately after electroporation, 4 ml of pre-warmed recovery medium was added to the cells, and the mixture was incubated at 37°C for 1 hour. Following recovery, 4 µl of the transformed cells was diluted to 1 ml with LB medium. A 100 µl aliquot of this dilution was spread on an LB plate containing the appropriate antibiotic. The number of resulting colonies was used to estimate the library diversity, which was calculated to be approximately 106 variants. The remaining transformation mixture was spread across four large (15 cm) LB plates containing antibiotics. After overnight incubation, 10 ml of LB liquid medium was added to the surface of each plate, and the bacterial colonies were collected by gently scraping. This process was repeated two to three times, and the collected bacterial suspension was pooled into a single tube. The plasmid library was purified using a Maxi Prep kit, following the manufacturer’s protocol.

### Viral Transduction

HEK293T was cultured in D-MEM (Wako) containing 10% fetal bovine serum (FBS) (biowest). HEK293T was authenticated by short-tandem repeat analyses and tested for mycoplasma contamination in our laboratory. All cell lines were cultured at 37 ℃ and 5 % CO2. Lentiviruses and retroviruses were produced by transient transfection in 293T cells using the calcium-phosphate method (Kitamura et al., 2003). Mixture of plasmids (3 µg of VSVG and 10 µg of PAX2 for lentivirus together with 12 µg virus vectors) (25 ml), 2.5 M CaCl2 (50 ml) and filtered water (425 ml) to 2×HeBS (500 ml) was added to 293T cells in a 10 cm dish (2.0×10^6^ cells/dish) for 18 hours. After 24 hours, we removed medium by aspiration and added 5 ml of fresh DMEM(Wako)/10% FBS (biowest). Virus-containing medium was collected 48 hours after the transfection. Supernatant was retrieved and centrifuged for 3 hrs at 3,000 g with LT-HSC. After centrifuge, remove medium by aspiration and add fresh 200 µl of StemSpan™ SFEM (STEMCELL Technologies Inc.).

### HSC Culture

HSC culture media consists of StemSpan™ SFEM (STEMCELL Technologies Inc.) supplemented with 50 ng/ml recombinant mouse SCF (R&D systems), 50 ng/ml recombinant human TPO (R&D systems).

### Bone Marrow Transplantation

The following barcode infected hematopoietic stem cells were transplanted to lethally irradiated mice (9.5 Gy, CD45.2) by tail vein injection with 2 ×10^5^ whole BM cells from C57BL/6-Ly5.2 (Ly5.2). For the secondary transplantation assay, 1/100 of whole BM cells from primary recipient mice 16 weeks after transplantation were transplanted into lethally irradiated (9.5 Gy) CD45.2 mice.

### Barcoding sequence

BM cells and spleen cells were collected from 16 weeks after transplantation and incubated with Lineage antibodies CD41-APC (clone MWReg30) (1:400), CD71-PE (clone RI7217) (1:400), CD19-APCcy7 (clone 6D5) (1:400), CD3e-BV785 (clone 145-2C11) (1:400), CD11b-PEcy7 (clone M1/70) (1:400) in 1ml for 15 minutes at 4 ℃. Following the addition of 1 mL of PBS and centrifugation for 5 minutes, the supernatant was discarded. The cells were resuspended in 4 mL of PBS and sorted for the 10^5^ BFP+CD41+ cells and BFP+CD71+ cells from BM and BFP+CD11b+ cells, BFP+CD19+ cells and BFP+CD3e+ cells from spleen. DNA was extracted using Nucleospin Blood (Takara). Barcode locus was amplified by 2 stages-PCR. In 1^st^ stage, PCR reactions were constructed 10 ml of genomic DNA and 0.25 mM Primer 1 (GACCTCCCTAGCAAACTGGGGCACAAG), and 0.25 mM Primer 2 (AGCATGCCTGCTATTGTCTTC), and amplified using KOD Fx Neo (Toyobo) according to the following PCR protocol: (1) 94℃ for 2 min, (2) 98 ℃ for 10 s, 50 ℃ for 30 s then 70 ℃ for 60 s (23 cycles). In 2^nd^ stage, PCR reactions were constructed 10 ml of 1^st^ PCR product and 0.25 mM Primer 1 (AATGATACGGCGACCACCGAGATCTACACGATCGGAAGAGCACACGTCTG AACTCCAGTCAC [Truseq ID 12, 6, 14, 10, 9, 1, 23, 13, 5, 4, 7 and 11] GACCTCCCTAGCA AACTGGGGCACAAG), and 0.25 mM Primer 2 (CAAGCAGAAGACGGCATACGAGATAGC ATGCCTGCTATTGTCTTC), and amplified using KOD Fx Neo (Toyobo) according to the following PCR protocol: (1) 94℃ for 2 min, (2) 98 ℃ for 10 s, 50 ℃ for 30 s then 70 ℃ for 60 s (23 cycles). Fragments of length 386 were selected by the electrophoresis and then extracted using Wizard SV Gel and PCR Clean-Up System (Promega). Barcode libraries were sequenced by HiSeq 2500.

### Barcode data process

The barcode information was extracted using bartender (Zhao et al., 2018). The sequences extracted by bartender were clustered, and the sequence with the largest number of reads in one cluster was evaluated as the barcode that the clone had, and the frequency of the clone was calculated from the percentage of reads. In addition, the following two treatments were performed to reduce the effects of sequencing, PCR errors, and Truseq-ID sequencing errors. Barcodes with less than 10 reads were considered to have no reads. Specimens with only 1/100th of the number of reads of all specimens were considered to have no detectable barcodes. K-means clustering was performed using ComplexHeatmap 1(Gu, 2022; Gu et al., 2016). Clusters were annotated based on their lineage output patterns, and annotated clusters were used for downstream analysis.

### Single cell RNA-seq

scRNA-seq libraries were constructed using the Chromium Single Cell 3′v3.1 Reagent Kit (10x Genomics, PN-1000269, PN-1000127, and PN-1000213/2000240) according to the manufacturer’s protocol. Briefly, the post-sorting sample volume was reduced. Cells were then loaded into each channel with a target output of approximately 5,000 cells. Single cells were encapsulated into emulsion droplets using the Chromium Controller (10x Genomics). Reverse transcription, fragmentation, and indexing PCR were performed in a thermal cycler. cDNA and final libraries were purified using AMPureXP beads (Beckman Coulter, A63881). A random DNA barcode sequence was amplified using the SI primer (10x Genomics, PN-2000095) and the Reverse primer (GAGTTCAGACGTGTGCTCTTCCGATCTGACCTCCCTAGCAAACTGGGGCACAAG) following this protocol: 98°C for 2 minutes, 10 cycles of (98°C for 10 seconds, 60°C for 30 seconds, and 68°C for 30 seconds), and 72°C for 10 minutes. Indexing PCR was then performed using the SI primer and NEBNext® Multiplex Oligos for Illumina® (Index Primers Set, NEB E7335S) following this protocol: 98°C for 2 minutes, 5 cycles of (98°C for 10 seconds, 60°C for 30 seconds, and 68°C for 30 seconds), and 72°C for 10 minutes. Amplified cDNA and final libraries were evaluated on an Agilent BioAnalyzer using a High Sensitivity DNA Kit (Agilent Technologies, 5067-5584 and 5067-5585). Individual libraries were diluted to 4nM and pooled for sequencing. The pools were sequenced using 150-cycle run kits (26bp Read1, 8bp Index1, and 90bp Read2) on the Nova-seq SP Sequencing System (Illumina).

### Single cell RNA-seq data process

The barcode information from scRNA-seq was extracted using bartender. The sequences extracted by bartender were clustered, and the sequence with the largest number of reads in one cluster was evaluated as the barcode that the clone had. The barcode with the highest number of reads in a single cell was evaluated to be the barcode with that cell. If the number of the reads were under 1,000, that cells evaluate as no barcode information. Single-cell RNA-seq data from three mouse samples were processed using Seurat(Hao et al., 2021). Count matrices were imported with Read10X and converted into Seurat objects, and cell-level metadata were added from external annotation files. Datasets were merged and quality-controlled by filtering cells with 300–10,000 detected genes and <5% mitochondrial transcripts. Data were log-normalized, highly variable genes were identified using the VST method, and expression values were scaled. Principal component analysis was performed on variable genes, followed by batch correction across samples using Harmony(Korsunsky et al., 2019), with sample identity as the batch variable. Harmony embeddings were used for neighborhood graph construction, clustering, and UMAP visualization (first 10 dimensions). Clusters were identified using a graph-based method and manually annotated. MolO scores were calculated using the following a publicly available pipeline (https://github.com/fionahamey/hscScore)(Hamey and Göttgens, 2019). Briefly, MolO scores were inferred by applying a pretrained hscScore regression model to log-transformed, total-count–normalized expression values derived from our scRNA-seq data.

### Statistics and reproducibility

Statistical analyses were performed by the unpaired and two-tailed Student’s t-test, analysis of variance (ANOVA) with Dunnett’s or Turkey’s post hoc test after testing for normal distribution and equal variance or Fisher’s exact test. GraphPad Prism 9.1.0 was used for these statistical analyses. The numbers of samples were presented as n in the figure legends. All data are presented as mean ± SEM in the figures. Significance levels were set at p* < 0.05, and p** < 0.01.

