## Supplementary material for "DNA barcoding–based paired daughter cell analysis reveals division preferences of hematopoietic stem cells": Figure S1-4

### FigureS1

A

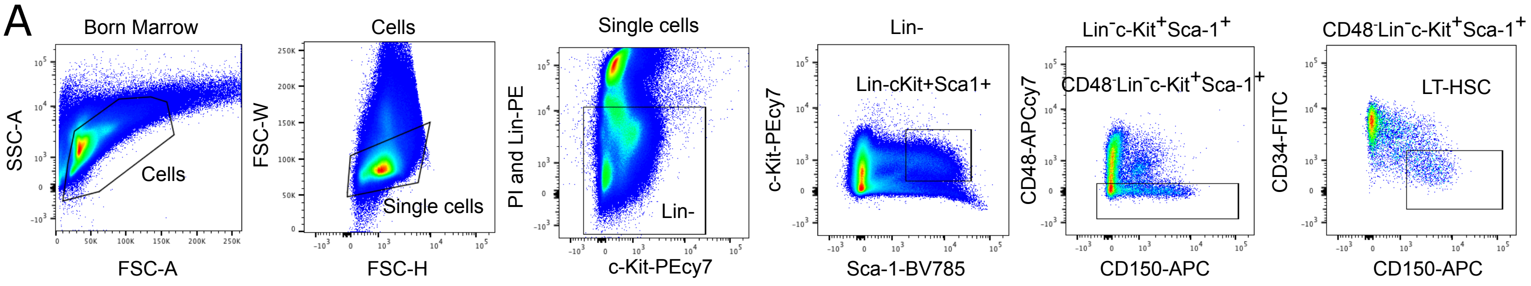


B

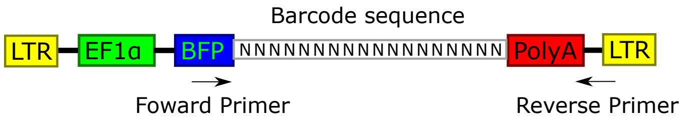


C

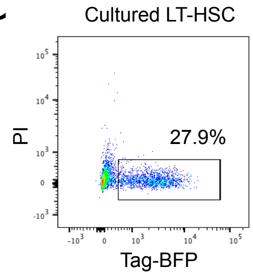


| | transduction efficiency |
|---|---|
| Ex1 | 27.9% |
| Ex2 | 24.6% |
| Ex3 | 35.1% |

D

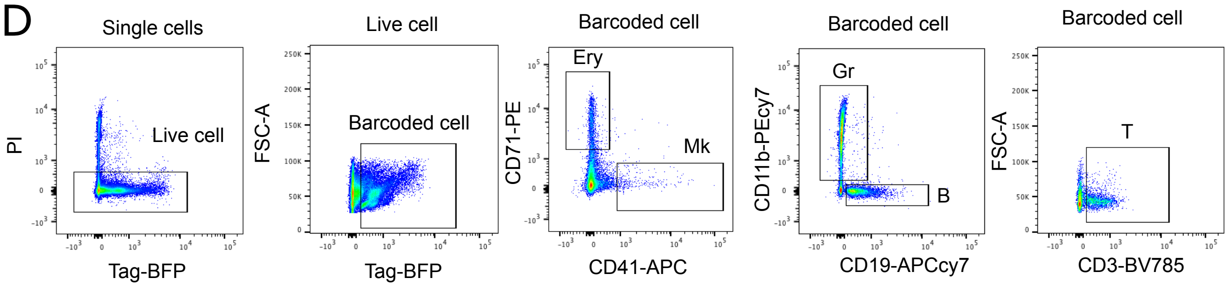


E

The proportion of barcoded cells in BM
16weeks after transplantation

| | mouse1 | mouse2 |
|---|---|---|
| Ex1 | 13.8% | 30.0% |
| Ex2 | 15.0% | 13.2% |
| Ex3 | 39.2% | 33.1% |

F

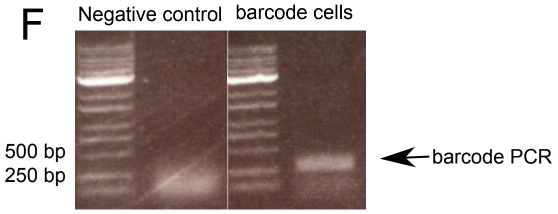

**Figure S1 | Isolation of LT-HSCs, barcode construct design, and validation of barcode transduction and detection.** **A**, Flow cytometry gating strategy for the isolation of HSCs from mouse bone marrow. **B**, Schematic of the lentiviral barcode construct. **C**, Representative flow cytometry plots showing BFP expression in cultured LT-HSCs after lentiviral transduction. Infection efficiency in independent experiments is indicated. **D**, Flow cytometric gating strategy for lineage analysis of barcoded cells. Live $BFP^+$ cells were analyzed for erythroid ($CD71^+$), megakaryocyte ($CD41^+$), granulocyte ($CD11b^+$), B cell ($CD19^+$), and T cell ($CD3^+$) markers. **E**, Quantification of the proportion of barcoded cells in peripheral blood at 4 and 16 weeks after transplantation in two independent experiments and two recipient mice. **F**, Agarose gel electrophoresis showing PCR amplification of barcode sequences. Negative control demonstrates specificity of barcode detection.

### Figure S2

A

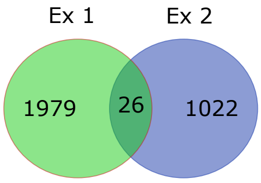


B

| | Pair | All barcodes |
|---|---|---|
| Ex1 | 405 | 1979 |
| Ex2 | 71 | 1022 |

C

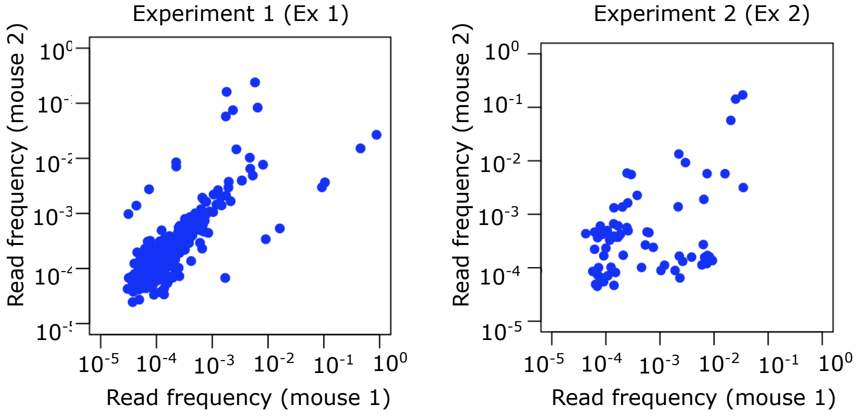


D

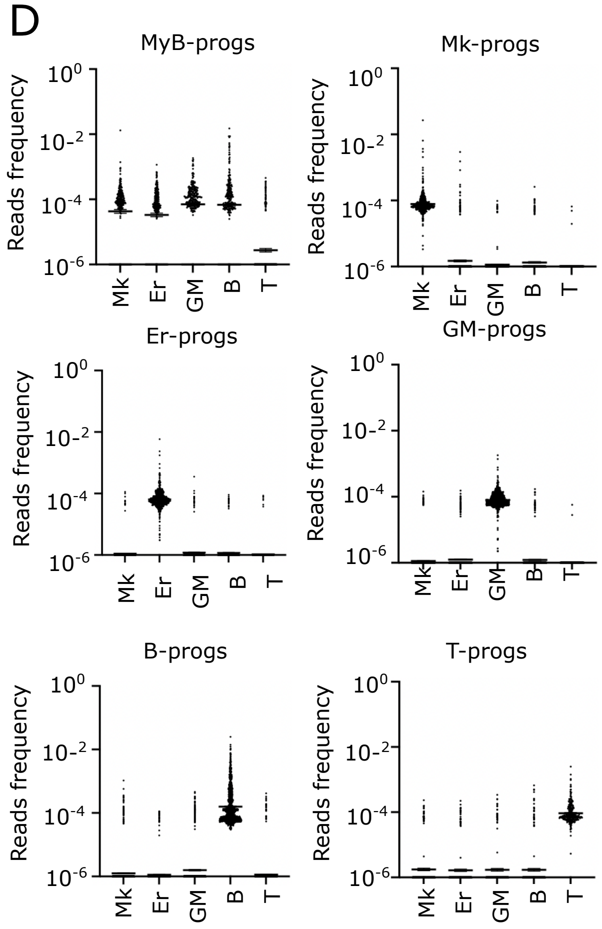


E

| | Stem | Prog |
|---|---|---|
| Stem | 50 ** $P<0.0001$ | 173 |
| Prog | | 3031 |

**Figure S2 | Reproducibility of paired daughter clone detection and lineage classification across independent experiments.** **A**, Venn diagrams showing overlap of detected daughter clones between independent experiments. **B**, Summary table of barcode detection outcomes in three independent experiments (Ex1–Ex3). The number of daughter pairs detected in both recipient mice (Pair), as well as all detected barcodes, are shown for each experiment. **C**, Scatter plots showing barcode read frequencies in recipient mouse 1 versus mouse 2 for Experiments 1–3. Each dot represents a barcode detected in both mice. Axes are displayed on a log scale. **D**, Dot plot showing the lineage output of progenitor-restricted clusters, including MyB-progs, Mk-progs, Er-progs, GM-progs, and B-progs. Each dot represents a daughter clone, and the y-axis indicates normalized barcode read frequency (log scale). These profiles confirm lineage-restricted output patterns used for cluster annotation in Figure 2. Statistical significance was assessed by one-way ANOVA followed by Tukey's multiple comparisons test (p values indicated). *$p < 0.05$ and **$p < 0.01$. n.s., not significant.

### Figure S3

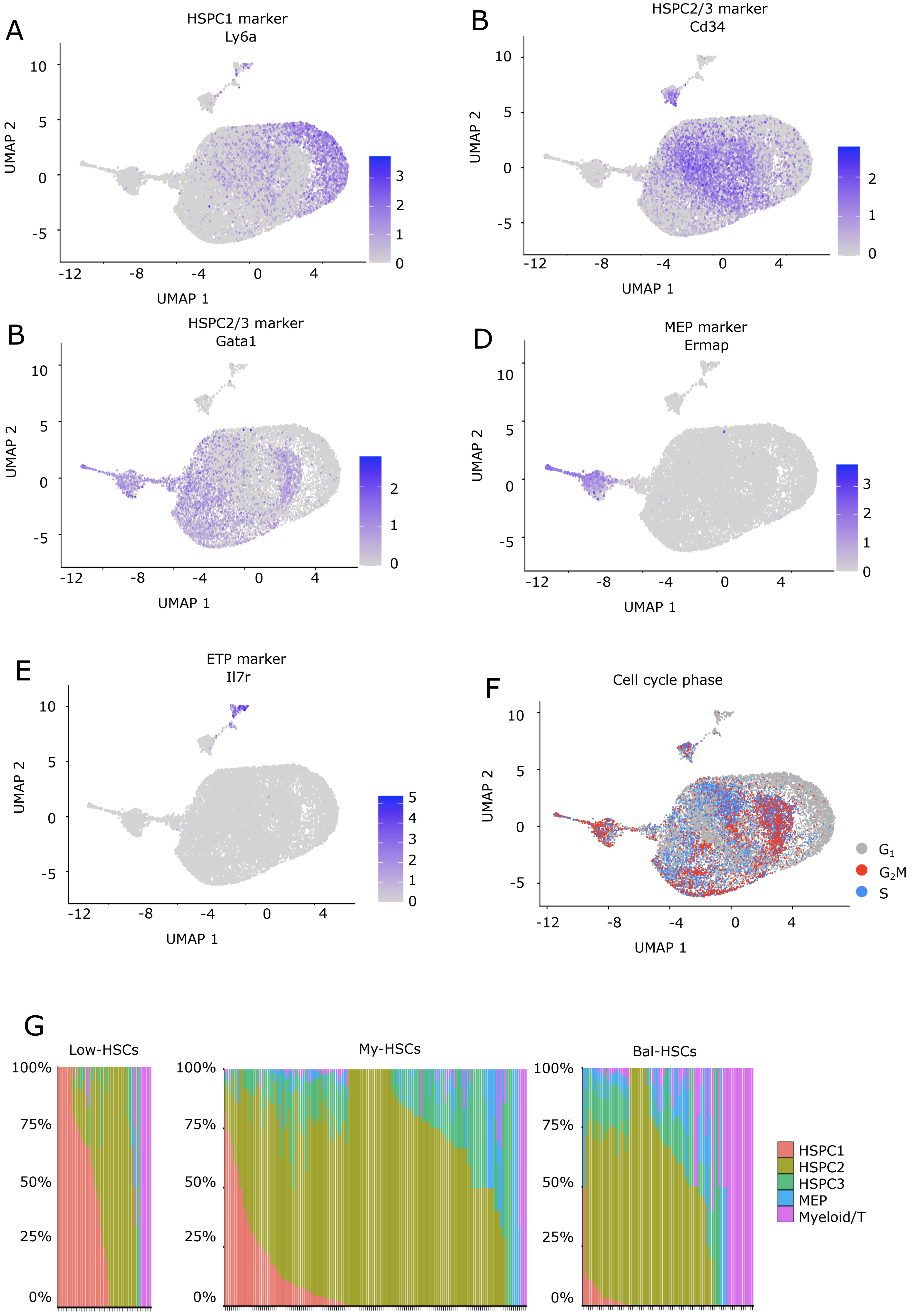
A
HSPC1 marker
Ly6a
B
HSPC2/3 marker
Cd34
B
HSPC2/3 marker
Gata1
D
MEP marker
Ermap
E
ETP marker
Il7r
F
Cell cycle phase
G1
G2M
S
UMAP 1
UMAP 2
G
Low-HSCs
My-HSCs
Bal-HSCs
100%
75%
50%
25%
0%
HSPC1
HSPC2
HSPC3
MEP
Myeloid/T

**Figure S3 | Marker gene expression and cell cycle annotation of transcriptional clusters.** **A–E**, Feature plots showing expression of representative cluster-specific marker genes overlaid on the UMAP projection. **F**, Cell cycle phase annotation projected onto the UMAP embedding. **G**, Proportional distribution of transcriptional clusters across daughter clones. Bars represent the percentage of transcriptional clusters within each clone.

### FigureS4

A

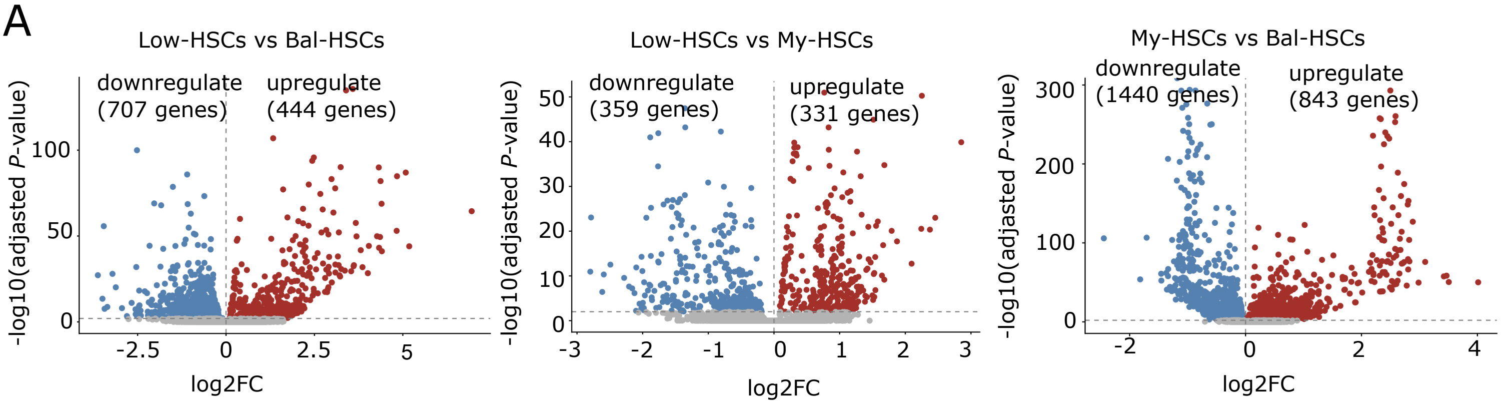

Low-HSCs vs Bal-HSCs
downregulate (707 genes)
upregulate (444 genes)
-log10(adjasted P-value)
log2FC
Low-HSCs vs My-HSCs
downregulate (359 genes)
upregulate (331 genes)
-log10(adjasted P-value)
log2FC
My-HSCs vs Bal-HSCs
downregulate (1440 genes)
upregulate (843 genes)
-log10(adjasted P-value)
log2FC

**Figure S4 | Differential gene expression, secondary transplantation outcomes, and transcriptional cluster distribution across HSC subsets. A**, Volcano plots showing differential gene expression between HSC subsets, restricted to cells in HSC1-3. Left: Low-HSCs versus Bal-HSCs. Middle: My-HSCs versus Bal-HSCs. Right: Low-HSCs versus My-HSCs. The x-axis represents log2 fold change (log2FC), and the y-axis represents -log10 adjusted p value. Numbers of significantly upregulated and downregulated genes are indicated in each comparison.
